# NBCn1 interacts with DYNLL1 and regulates ciliary length and SUFU localization to control Sonic hedgehog signaling

**DOI:** 10.64898/2026.03.26.714467

**Authors:** Mohamed Chamlali, Louise Bødstrup Rosenkrantz, Emma Sloth Frandsen, Renée Patungan, Esben Lorentzen, Stine Falsig Pedersen, Lotte B. Pedersen

**Affiliations:** Department of Biology, University of Copenhagen, Denmark; Department of Molecular Biology and Genetics, Aarhus University, Denmark

**Keywords:** NBCn1, primary cilia, DYNLL1, VPS45, intraflagellar transport, Sonic hedgehog, SUFU, TMEM216

## Abstract

Primary cilia integrate morphogen signaling with cellular physiology, yet how ion transporters contribute to ciliary organization and function remains poorly understood. Here, we identify the sodium-bicarbonate cotransporter NBCn1 (SLC4A7) as a previously unrecognized ciliary membrane component essential for regulating ciliary length and Sonic hedgehog (Shh) signaling. NBCn1 localizes to primary cilia through distinct N⍰and C⍰terminal targeting motifs and inhibition of dynein activity or loss of DLG1, a known NBCn1 interacting protein, augments its ciliary localization. Loss of NBCn1 shortens cilia without altering ciliation frequency or deciliation kinetics, indicating a selective role in ciliary elongation control rather than in ciliogenesis initiation or maintenance. NBCn1 deficiency or -inhibition leads to ciliary enrichment of SUFU even at basal level and markedly attenuates GLI1 transcriptional activation, despite intact SMO ciliary entry. Structural modeling and co⍰immunoprecipitation reveal that NBCn1 interacts with DYNLL1, VPS45, and the transition⍰zone protein TMEM216, suggesting that NBCn1 modulates ciliary length and Shh signaling through interactions with these proteins. Together, these findings uncover NBCn1 as a central regulator of ciliary length and Shh signaling, highlighting ion transport as a critical and underappreciated determinant of ciliary signaling competence.

**Significance statement:** Primary cilia are essential cellular signaling hubs, yet the contribution of ion transporters and pH to ciliary architecture and function has remained unclear. This study identifies the bicarbonate transporter NBCn1 as a previously unrecognized regulator of ciliary length and Sonic hedgehog (Shh) signaling. We show that NBCn1 is actively trafficked into cilia through defined N⍰and C⍰terminal determinants, is exported from cilia via retrograde intraflagellar transport (IFT), and interacts directly with IFT dynein component DYNLL1. Loss of NBCn1 selectively shortens cilia while leaving ciliogenesis frequency and deciliation dynamics intact. NBCn1 deficiency disrupts the ciliary localization of the key Shh component SUFU, leading to impaired GLI transcriptional responses. By revealing how ion transport intersects with IFT and morphogen signaling, these findings establish NBCn1 as a central integrator of ciliary microenvironmental regulation and developmental signaling output, highlighting pH regulatory ion transport as a critical and underappreciated determinant of ciliary function.

## Introduction

The sodium–bicarbonate cotransporter NBCn1 (SLC4A7) is a principal regulator of net acid extrusion and plays essential roles in intracellular pH control, cell proliferation, and growth [1, 2]. NBCn1 is broadly expressed in epithelial and non-epithelial tissues [2, 3], and altered NBCn1 activity contributes to major human diseases, including hypertension and cancer [4–6]. NBCn1 plays important roles in kidney physiology [2] and we previously found that NBCn1 is targeted to the basolateral surface of polarized kidney epithelial cells (MDCK II) through interactions mediated by its cytoplasmic C terminus. This region binds protein complexes involved in membrane trafficking, including endocytic regulators, the retromer and WASH complexes, and components of the Scribble polarity machinery [7].

The Scribble complex, composed of SCRIB, DLG1 and LGL in vertebrates, localizes to the basolateral domain of epithelial cells and is required for establishing and maintaining apical–basal polarity [8]. Consistent with biochemical data showing that NBCn1 associates with multiple Scribble components [7], NBCn1 also interacts with the DLG1 homolog DLG4 in rat brain lysates [9]. Using co immunoprecipitation and proximity ligation assays, we recently confirmed a physical interaction between NBCn1 and DLG1 and identified molecular elements required for NBCn1 accumulation at the plasma membrane [10]. During these studies, we observed that NBCn1 is strongly enriched in the primary cilium of several polarized epithelial cell types [10]. In addition, we demonstrated that DLG1 plays a key role in regulating ciliary length and protein content in kidney epithelial cells. Specifically, we showed that DLG1 functions together with retromer-associated protein SDCCAG3 and IFT20 to promote trafficking of polycystin-2 to the primary cilium [11]. However, despite its physical association with DLG1 as well as with retromer components [7], the mechanisms that mediate NBCn1 ciliary localization, and the functional role of NBCn1 within the cilium, remain unknown.

Primary cilia are microtubule-based organelles that extend from the surface of many vertebrate cell types, including kidney epithelial cells, and are essential for regulating multiple signaling pathways during development and in adult tissue physiology. Defects in ciliary structure or function disrupt signaling and underlie a large group of human disorders collectively termed ciliopathies, which affect a wide range of organs such as kidney, brain, and heart [12]. Examples include autosomal dominant polycystic kidney disease, caused by defects in the polycystin-1/2 ciliary receptor–channel complex [13], and Bardet–Biedl syndrome (BBS), which results from mutations in the BBSome trafficking complex and is characterized by renal, metabolic, and neurological abnormalities [14].

The primary cilium consists of an axonemal microtubule scaffold surrounded by a membrane enriched in distinct receptors, ion channels, and signaling lipids. One of the most extensively characterized pathways operating within this compartment is the Sonic hedgehog (Shh) pathway, which is an essential regulator of multiple processes in development and tissue homeostasis, and dysregulation of which plays a critical role in several cancers [15, 16]. The primary cilium serves as the central organizing hub for vertebrate Shh signaling by spatially coordinating the trafficking and processing of pathway components. In the absence of ligand, PTCH1 localizes to the cilium to prevent Smoothened (SMO) ciliary accumulation, enabling SUFU to sequester GLI transcription factors and promote formation of GLI repressors. Upon Shh binding, PTCH1 exits and SMO accumulates in the cilium, triggering ciliary tip events that weaken SUFU–GLI interactions and allow full-length GLI proteins to convert into activators that drive transcriptional responses. Through this ligand-dependent transition between SUFU-mediated repression and cilium-dependent activation, the primary cilium ensures precise temporal and spatial control of Shh signaling [12].

Ciliary signaling requires tight control of the protein and lipid content of the ciliary membrane. This composition is dynamically regulated through coordinated import and export processes, including intraflagellar transport (IFT) and BBSome-mediated trafficking, selective gating at the transition zone, and removal of proteins via endocytosis or extracellular vesicle release [17, 18]. However, the mechanisms by which cilia precisely adjust their molecular composition to tune signaling outputs remain incompletely defined.

Here, we identify sequence elements that govern NBCn1 targeting to the primary cilium and show that these motifs, together with motor⍰and adaptor⍰dependent trafficking pathways, position NBCn1 as an active component of the ciliary membrane. We further demonstrate that NBCn1 is required to establish normal ciliary length, regulate the ciliary localization of Shh pathway component SUFU in the basal state, and enable full transcriptional activation of Shh target gene expression in response to pathway activation. Together, these findings uncover an unexpected link between bicarbonate transport, ciliary architecture, and morphogen signaling, revealing NBCn1 as a previously unrecognized regulator of ciliary function.

## Results

### Ciliary targeting sequences in NBCn1

We previously showed that NBCn1 (encoded by *Slc4a7*) localizes to the primary cilium of different mammalian epithelial cell types [10], but how NBCn1 is targeted to, and may function at, the primary cilium is unknown. To define the molecular determinants responsible for ciliary localization of NBCn1, we expressed a series of GFP-tagged deletion constructs of NBCn1, which either lack the entire N- or C-terminal cytosolic regions (denoted ΔN- and ΔC⍰terminus, respectively) or specific amino acid residues within these regions (Δ1-98, Δ1133–1136, Δ1136–1139, Δ1139–1254; Fig.□1A) [7, 10], in ciliated mouse inner medullary collecting duct 3 (IMCD3) cells; we then examined their ciliary localization by live-cell confocal microscopy using SiR-tubulin to stain cilia and centrioles. Quantitative fluorescence imaging showed that removal of the entire N- or C-terminal cytosolic regions significantly impaired ciliary targeting of NBCn1 by 89 and 95%, respectively, as compared to the full-length, wild-type (WT) protein (Fig.□1B-D). Furthermore, deletion of residues 1133-1136 (Δ1133–1136) within the C-terminal cytosolic tail led to a significant reduction of NBCn1 ciliary localization whereas ciliary localization of the Δ1-98, Δ1136–1139, and Δ1139–1254 fusion proteins was similar compared to WT NBCn1 (Fig. 1B-D). These data suggest that NBCn1 ciliary targeting or retention requires elements located within residues 99-611 in the N-terminal region, as well as residues 1133-1136 in the C-terminal region.

**Figure 1:**
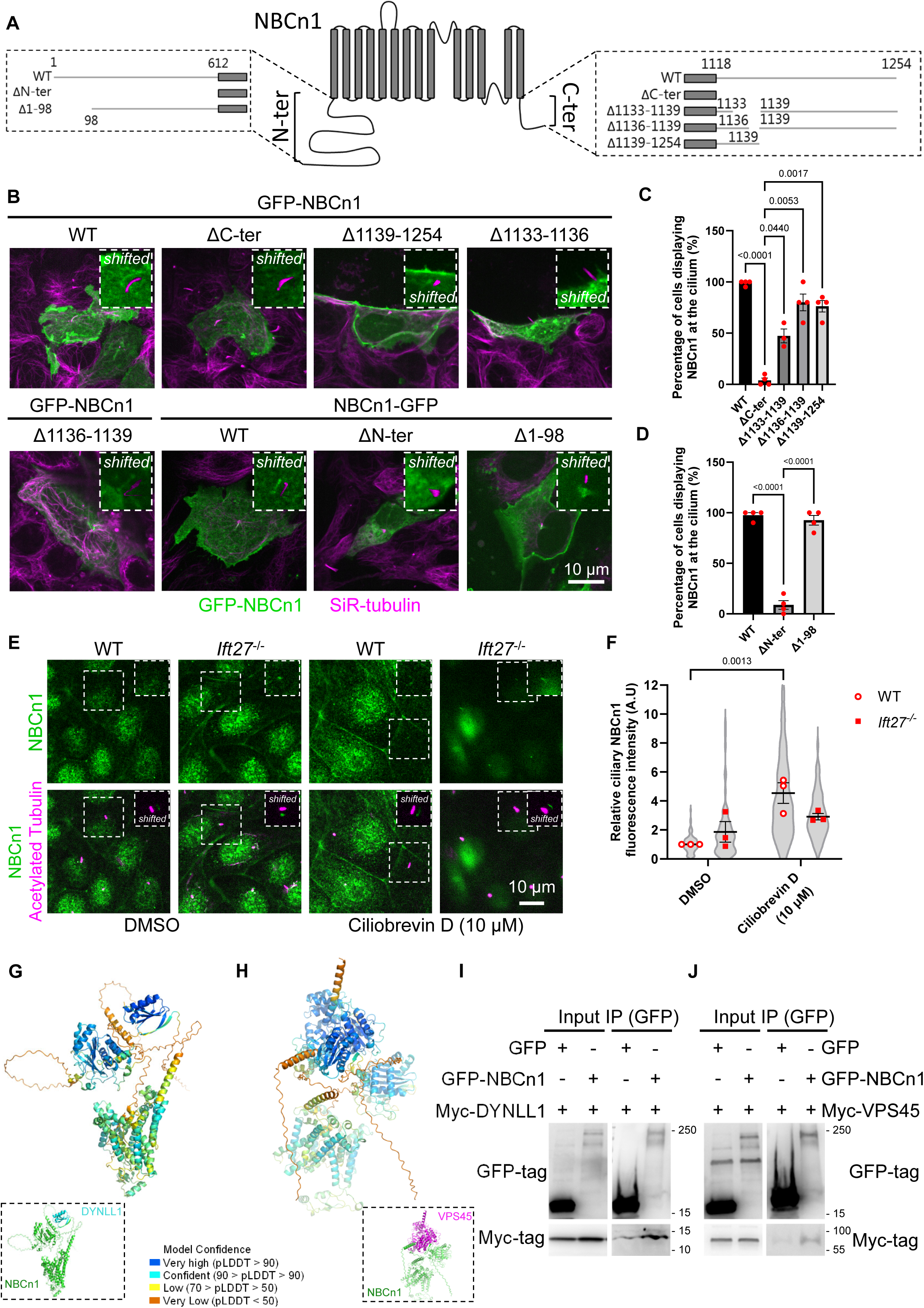
Targeting mechanisms of NBCn1 to the primary cilium. **A.** Overview of the C- and N-terminal cytosolic regions of NBCn1 and constructs used in this study. **B.** Confocal images of cells expressing GFP-tagged NBCn1 full-length/WT or the indicated deletion constructs. SiR-tubulin (magenta) stains primary cilia. Insets show magnified views of the cilium (“shifted” panels). Scale bar, 10 µm. **C.** Percentage of cilia displaying NBCn1 (GFP-tag in N-terminus) enrichment normalized to WT. Each red point represents one biological replicate; bars show mean ± SEM; statistical significance was determined by one-way ANOVA with multiple comparison testing (N=3-4). **D.** Percentage of cilia with NBCn1 (GFP-tag in C-terminus) enrichment normalized to WT. Each red point represents one biological replicate; bars show mean ± SEM; statistical significance was determined by one-way ANOVA with multiple comparison testing (N=3-4). **E.** Representative confocal images of WT and *Ift27*^-/-^ cells before and after treatment with 10 μM Ciliobrevin D stained for NBCn1 (green), acetylated α-tubulin (magenta) to visualize primary cilia. Scale bar, 10 µm. **F.** IFM analysis of ciliated IMCD3 cell lines showing comparative ciliary NBCn1 staining in WT and *Ift27*^-/-^ cells before and after treatment with 10 μM Ciliobrevin D. Each red point represents one biological replicate; grey violin plots represent the technical replicates; bars indicate mean ± SEM (N= 3, WT DMSO n= 107, *Ift27*^-/-^DMSO n= 81, WT Ciliobrevin D n= 75, *Ift27*^-/-^ Ciliobrevin D n= 68); statistical significance was determined by two-way ANOVA with multiple comparison testing. **G.** AlphaFold3-predicted structure of the NBCn1–DYNLL1 complex, colored by pLDDT confidence score. The predicted interface involves residues 44–54 of NBCn1 in the N-terminal cytoplasmic domain and the canonical cargo-binding groove of DYNLL1 (residues 60–89). See Fig. S1E for PAE matrix. **H.** AlphaFold3-predicted structure of the NBCn1–VPS45 complex, colored by pLDDT confidence score. The predicted interface involves an overlapping region of NBCn1 (residues 45–51) and residues 119–126 of the VPS45 SEC1/MUNC18-like domain. See Fig. S1F for PAE matrix. **I.** IP of HEK293T cells co-expressing the indicated GFP-tagged NBCn1 fusion protein or GFP alone and Myc-DYNLL1. Cell lysates were subjected to anti-GFP IP, and pellet and input samples were analyzed by SDS-PAGE and western blotting with anti-GFP or anti-Myc. Molecular mass markers are shown in kDa to the right. **J.** IP of HEK293T cells co-expressing the indicated GFP-tagged NBCn1 fusion protein or GFP alone and Myc-VPS45. Cell lysates were subjected to anti-GFP IP, and pellet and input samples were analyzed by SDS-PAGE and western blotting with anti-GFP or anti-Myc antibodies. Molecular mass markers are shown in kDa to the right.

### Ciliary localization of NBCn1 is negatively regulated by DLG1 and retrograde IFT

Previous work from our labs has shown that the Scribble polarity complex protein DLG1 regulates the length and composition of primary cilia in mouse kidney epithelial cells [11], and that DLG1 interacts physically with NBCn1 [10]. We therefore tested if ciliary localization of NBCn1 requires DLG1 by comparing the ciliary localization of full-length, GFP-tagged NBCn1 in WT and our previously described *Dlg1*^-/-^ IMCD3 cells [11]. Quantitative live-cell confocal imaging analysis of cells expressing GFP⍰tagged NBCn1 and with cilia visualized by SiR-tubulin staining revealed that the *Dlg1*^-/-^ cells displayed a significant (38%) increase in relative NBCn1 ciliary levels as compared to WT (Fig.□S1A, B), suggesting that DLG1 inhibits ciliary targeting of NBCn1 and/or promotes its retrieval from the organelle.

To further explore the mechanisms involved in regulating ciliary NBCn1 targeting and homeostasis, we next focused on components of the IFT machinery, in particular the Rab-like GTPase IFT27, which is part of the IFT-B complex and thought to promote ciliary export of select membrane proteins by linking the BBSome to retrograde IFT trains [19–21]. Interestingly, our previous work showed that IFT27 interacted with the NBCn1 C-terminal tail in a pull-down assay [7, 10], suggesting a potential role for IFT27 in ciliary NBCn1 homeostasis. When we assessed the ciliary localization of GFP-tagged NBCn1 in WT and *Ift27*^-/-^ IMCD3 cells [22] by quantitative live-cell confocal microscopy, NBCn1 appeared to accumulate within cilia of the *Ift27*^-/-^ cells although the difference was not statistically significant compared to WT cells (Fig. 1E, F). In contrast, acute inhibition of dynein and therefore of retrograde IFT with 10 μM Ciliobrevin D [23], which caused significant ciliary accumulation of the IFT-B component IFT88 as expected (Fig.□S1C, D), led to increased relative ciliary levels of NBCn1 in WT and *Ift27*^-/-^ cells by 4.5- and 1.56-fold, respectively (Fig.1E, F). This suggests a role for retrograde IFT in mediating ciliary export of NBCn1.

Next, we asked how NBCn1 might associate with the retrograde IFT machinery during export from the cilium. AlphaFold3 structure predictions [24] for pairwise complexes of full-length mouse NBCn1 against the human ciliary proteome [25] identified dynein light chain LC8-type 1 (DYNLL1; iPTM 0.64) and vacuolar protein sorting-associated protein 45 (VPS45; iPTM 0.61) as predicted interaction partners (Fig. 1G, H; Fig. S1E, F). DYNLL1 is a component of the cytoplasmic dynein 1 and 2 complexes with a well-characterized role in retrograde IFT [26, 27], while VPS45 regulates vesicular trafficking and was shown to control ciliary homeostasis of polycystin-2 in *C. elegans* sensory neurons [28]. Both predicted interfaces map to residues 44–54 of NBCn1 in the N-terminal cytoplasmic domain: DYNLL1 engages the canonical cargo-binding groove (residues 60–89) through a combination of hydrophobic and polar contacts including two predicted salt bridges, while VPS45 interacts via residues 119–126 within its SEC1/MUNC18-like domain. The overlapping binding sites suggest that these two interactions are mutually exclusive, potentially reflecting sequential trafficking steps. Co-immunoprecipitation confirmed both interactions: GFP-NBCn1, but not GFP alone, co-precipitated Myc-DYNLL1 (Fig. 1I) and Myc-VPS45 (Fig. 1J). Moreover, since residues 44–54 are dispensable for ciliary targeting (Fig. 1B, D), these data suggest that DYNLL1 and VPS45 mediate trafficking of NBCn1 out of, rather than into, the primary cilium.

### Loss or inhibition of NBCn1 leads to ciliary shortening

To investigate the function of NBCn1 at the primary cilium, we used CRISPR/Cas9 methodology to knock out (KO) the corresponding gene in IMCD3 cells. We verified two independent *Slc4a7*^-/-^ clones (Cl1 and Cl2) by INDEL sequencing analysis targeting the *Slc4a7* locus, confirming successful gene disruption (Fig.□2A); western blot analysis further verified the loss of NBCn1 protein in the two KO clones (Fig.□2B). To assess the impact of the loss of NBCn1 on intracellular pH (pH_i_), we monitored pH_i_ recovery following ammonium prepulse-mediated intracellular acidification under CO_2_/HCO_3-_ buffering conditions. Consistent with loss of net acid extrusion upon NBCn1 KO, both *Slc4a7*^-/-^ Cl1 and Cl2 exhibited an approximately 50% decrease in the rate of pH□ recovery (Fig.□2C, D). These data show that loss of NBCn1 results in reduced pH_i_ recovery capacity in IMCD3 cells, consistent with the key role of NBCn1 in pH_i_ regulation in most cell types including renal epithelial cells [2, 29].

Given that pH_e_ changes – which rapidly change pH_i_ in the same direction [30] – can induce ciliary disassembly [31] and potentially regulate ciliary length [32], we next examined how loss of NBCn1 affects ciliary homeostasis and length. Confocal fluorescence imaging of primary cilia revealed significant ciliary shortening in both *Slc4a7*^-/-^ clones compared with WT controls, both after 24 and 48□h of serum starvation to induce ciliogenesis (Fig.□2E, F). Specifically, cilia were 28 and 41% shorter in *Slc4a7*^-/-^ Cl1 and Cl2, respectively at 24 h, and 38 and 28% shorter in *Slc4a7*^-/-^ Cl1 and Cl2 at 48 h, compared to WT cells (Fig. 2E, F). In contrast, ciliation frequency was not affected by loss of NBCn1 (Fig. S2A, B), indicating that NBCn1 specifically promotes axoneme extension and is dispensable for initiating ciliogenesis. Supportively, treatment of WT IMCD3 cells with the NBCn1 inhibitor S0859 (20 μM) [33] before and during the onset of ciliogenesis (24□h) led to significant ciliary shortening (cilia length reduced by 39%) when compared to vehicle-treated controls (Fig.□2G, H). However, when S0859 was administered after cilia had already formed (24 h + 24 h), ciliary length remained similar between DMSO-treated and S0859⍰treated cells (Fig.□2G, H). Ciliation frequency was unaffected by S0859 treatment (Fig. S2C), further indicating that NBCn1 activity is required for ciliary elongation but not for initiation of ciliogenesis. Using similar approaches, we next asked whether NBCn1 contributes to cilium resorption in response to mitogenic stimulation. Specifically, we examined the kinetics of serum⍰induced deciliation in WT and *Slc4a7*^-/-^ cells using an established protocol (Fig.□S3A) [34], and found that ciliary disassembly rates were similar between WT and *Slc4a7*^-/-^ clones (Fig.□S3B-D). Thus, while NBCn1 is essential for ciliary elongation (Fig.□2, Fig. S2), its loss does not impair the ability of IMCD3 cells to disassemble primary cilia in response to serum stimulation.

**Figure 2:**
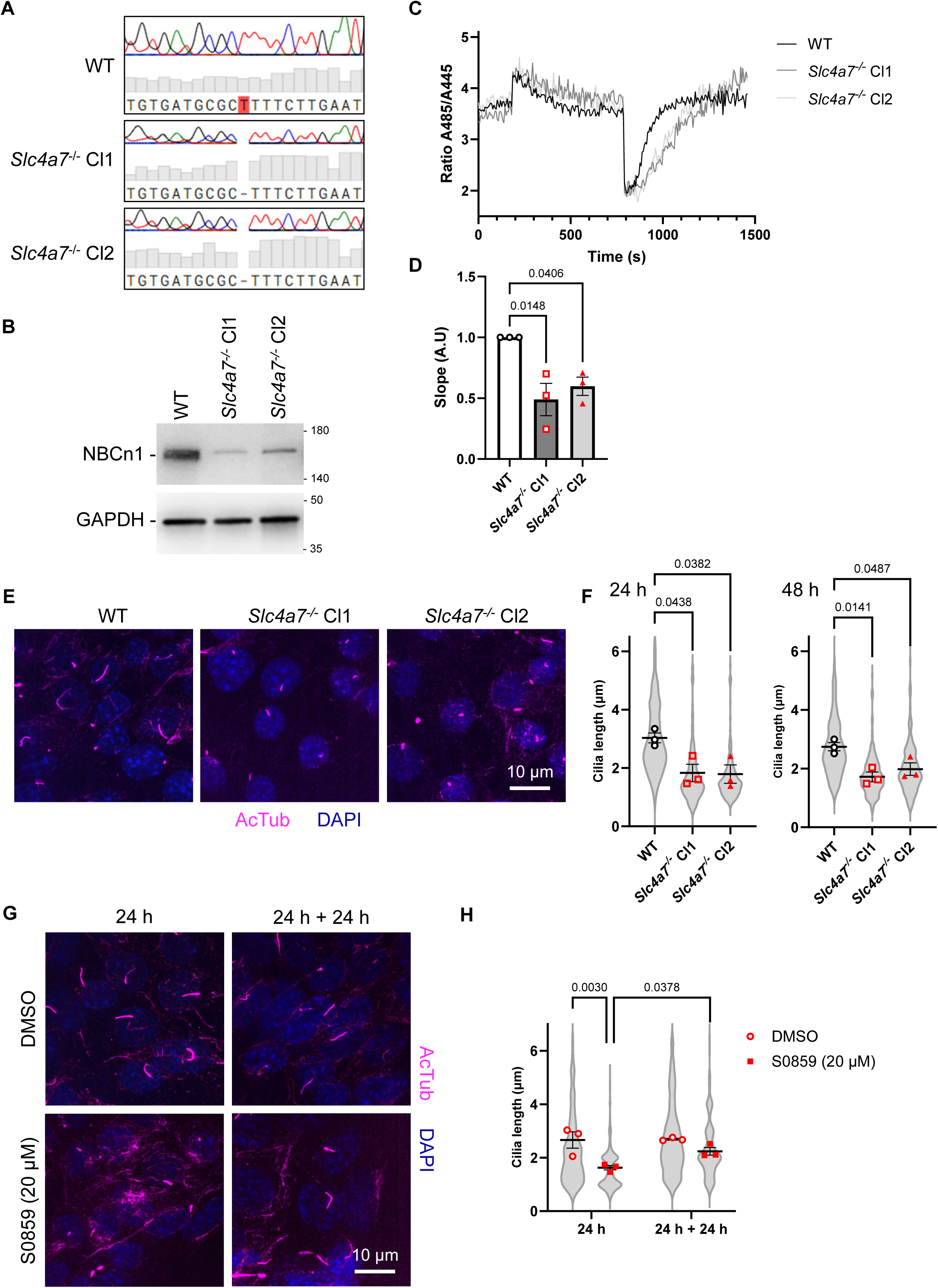
Loss of NBCn1 leads to ciliary shortening. **A.** INDEL analysis of the NBCn1 target locus in WT and *Slc4a7*^-/-^ IMCD3 cells. WT cells exhibit an unmodified sequence, while *Slc4a7*^-/-^ cells display insertions and/or deletions at the expected cut site, confirming CRISPR/Cas9-mediated gene knockout. **B.** Western blot analysis of total cell lysates of the indicated IMCD3 cells lines using antibodies against NBCn1 and GAPDH (used as loading control). Molecular mass markers are shown in kDa. **C.** pHi recovery measured after acidification by the ammonium prepulse technique in the presence of CO_2_ and bicarbonate buffer in WT versus *Slc4a7*^-/-^ IMCD3 cells. **D.** Quantification of the pHi recovery slope in WT versus *Slc4a7*^-/-^ IMCD3 cells. Each black/red point represents one biological replicate; bars indicate mean ± SEM (N= 3); statistical significance was determined by one-way ANOVA with multiple comparison testing. **E.** Representative confocal images of WT and *Slc4a7*^-/-^ cells (Cl1, Cl2) stained for acetylated α-tubulin (magenta) to visualize primary cilia and DAPI (blue) for nuclei. Scale bar, 10 µm. **F.** Quantification of ciliary length in WT and *Slc4a7*^-/-^ clones at 24 h and 48 h post starvation. Each black/red point represents one biological replicate; Grey violin plots represent the technical replicates; bars indicate mean ± SEM (N= 3, 24 h WT n= 168, 24 h *Slc4a7*^-/-^ Cl1 n= 153, 24 h *Slc4a7*^-/-^ Cl2 n= 173, 48 h WT n= 189, 48 h *Slc4a7*^-/-^ Cl1 n= 261, 48 h *Slc4a7*^-/-^ Cl2 n= 154); statistical significance was determined by one-way ANOVA with multiple comparison testing. **G.** Representative images of cells treated with DMSO or the NBCn1 inhibitor S0859 (20 μM) for 24 h and stained as above. Scale bar, 10 µm. **H.** Quantification of ciliary length after S0859 treatment. Cilia length was measured following S0859 treatment before (24 h) or after (24 h + 24 h) ciliogenesis. Each red point represents one biological replicate; grey violin plots represent the technical replicates; bars indicate mean ± SEM (N= 3, 24 h DMSO n= 300, 24 h S0859 n= 81, 24 h + 24 h DMSO n= 236, 24 h + 24 h S0859 n= 157); statistical significance was determined by two-way ANOVA with multiple comparison testing.

Together, these findings show that NBCn1 activity is essential for proper ciliary elongation in IMCD3 cells.

### Loss of NBCn1 affects Sonic hedgehog (Shh) pathway activation

One of the best characterized signaling pathways coordinated by the primary cilium is the Shh pathway [35], depicted schematically in Fig.□3A. Given the role of NBCn1 in promoting ciliary elongation, we therefore investigated whether it is required for Shh signaling. Ciliated WT and *Slc4a7*^-/-^ IMCD3 cells were treated with the SMO agonists purmorphamine (PMA) and SAG, or the PTCH1 ligand Shh, and RT⍰qPCR analysis of the Shh target gene *Gli1* was used as a readout of pathway activity. As expected, this analysis showed a significant increase in *Gli1* expression in WT cells treated with PMA, SAG or Shh, as compared to cells treated with vehicle alone (Fig. 3B; Fig. S4A). Notably, for both *Slc4a7*^-/-^ Cl1 and Cl2, the agonist-induced *Gli1* expression level was reduced by 79-90% compared to WT cells (Fig.□3B). Similar results, although not significant, were observed for *Gli1* transcript levels in response to ligand⍰driven pathway activation with Shh (Fig. S4A). These findings support the conclusion that NBCn1 is required for efficient Shh⍰dependent transcriptional output.

**Figure 3:**
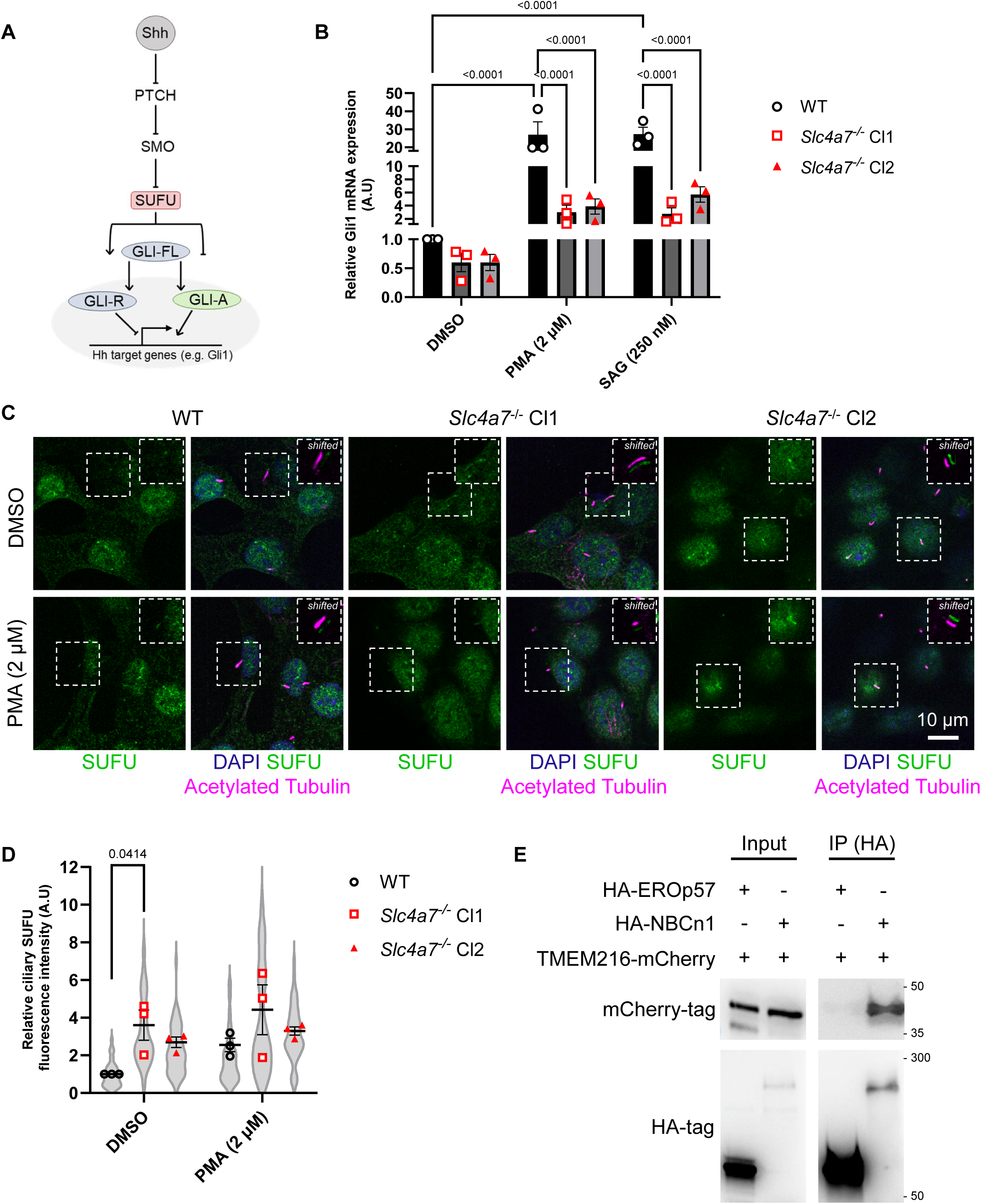
Loss of NBCn1 affects Sonic Hedgehog (Shh) signaling pathway activation. **A.** Schematic overview of the Shh signaling pathway [73]. **B.** RT-qPCR of Gli1 expression in WT and *Slc4a7*^-/-^ cells following stimulation with SMO agonists purmorphamine (PMA) and SMO agonist (SAG). Each black/red point represents one biological replicate; bars indicate mean ± SEM (N= 3); statistical significance was determined by two-way ANOVA with multiple comparison testing. **C.** Representative confocal images of WT and *Slc4a7*^-/-^ cells (Cl1, Cl2) stained for SUFU (green), acetylated α-tubulin (magenta) to visualize primary cilia and DAPI (blue) for nuclei. Scale bar, 10 µm. **D.** IFM analysis of ciliated IMCD3 cell lines showing comparative ciliary SUFU staining in WT and *Slc4a7*^-/-^ cells. Each black/red point represents one biological replicate; grey violin plots represent the technical replicates; bars indicate mean ± SEM (N= 3, DMSO WT n= 81, DMSO *Slc4a7*^-/-^ Cl1 n= 52, DMSO *Slc4a7*^-/-^ Cl2 n= 61, PMA WT n= 71, PMA *Slc4a7*^-/-^ Cl1 n= 69, PMA *Slc4a7*^-/-^ Cl2 n= 59); statistical significance was determined by two-way ANOVA with multiple comparison testing. **E.** IP of HEK293T cells co-expressing the indicated HA-tagged NBCn1 fusion protein or HA-EROp57 (acting as control protein) and TMEM216-mCherry. Cell lysates were subjected to anti-GFP IP, and pellet and input samples were analyzed by SDS-PAGE and western blotting with anti-HA or anti-mCherry antibodies. Molecular mass markers are shown in kDa to the right.

Because SMO ciliary accumulation is a pivotal early event in Shh signal propagation [36, 37], we next examined whether NBCn1 loss disrupts this process. Live-cell confocal imaging of GFP⍰SMO stably expressed in WT and *Slc4a7*^-/-^ parental IMCD3 cells revealed clear ciliary accumulation of SMO upon PMA or SAG treatment in both WT and *Slc4a7*^-/-^ cells (Fig.□S4B-E), indicating that SMO accumulates in cilia in response to agonist treatment regardless of NBCn1 status. Therefore, we next examined whether NBCn1 loss alters the ciliary localization of other key pathway components, such as Suppressor of Fused (SUFU) [36, 37]. Strikingly, confocal fluorescence imaging of SMO agonist (PMA) or vehicle (DMSO) treated cells revealed that SUFU accumulation in cilia was increased by 361% in *Slc4a7^-^*^/-^ cells compared to WT cells at basal level (DMSO) (Fig.□3C, D). In WT cells, PMA stimulation tended to increase ciliary SUFU levels compared to DMSO-treated cells although the difference was not statistically significant (Fig. 3C, D). For *Slc4a7*^-/-^cells, PMA stimulation did not further increase the ciliary SUFU level as compared to DMSO-treated cells. These findings indicate that NBCn1 limits ciliary accumulation of SUFU in the basal state (DMSO treatment), which could explain the failure of NBCn1-deficient cells to properly upregulate *Gli1* expression in response to pathway activation (Fig. 3B).

To explore the mechanism by which NBCn1 might regulate ciliary SUFU localization we used https://predictomes.org/, a database of AlphaFold-modeled protein-protein interactions [38, 39], to search for potential interactors of NBCn1 with links to SUFU. We identified a potential direct interaction between NBCn1 and TMEM216, a transmembrane component of the ciliary transition zone (the Meckel Gruber syndrome/Joubert syndrome (MKS/JBTS) complex) that was reported to modulate Shh signaling through its interaction with SUFU [40]. To experimentally test whether NBCn1 interacts physically with TMEM216, we performed a co⍰immunoprecipitation assay in HEK293T cells, which demonstrated that HA⍰tagged NBCn1, but not the control protein HA⍰EROp57, co⍰precipitated TMEM216⍰mCherry (Fig.□3E). These data suggest that NBCn1 binds physically to TMEM216, providing a potential mechanistic link between NBCn1⍰dependent regulation of ciliary SUFU localization and downstream activation of *Gli1* expression upon induction of Shh signaling.

## Discussion

### NBCn1 ciliary localization depends on discrete N⍰and C⍰terminal motifs and is regulated by DLG1 and retrograde IFT

We previously uncovered the key determinants of NBCn1 plasma membrane trafficking, and showed that NBCn1 localizes not only to the basolateral plasma membrane and cell–cell junctions of epithelial cells [7, 10], but also to primary cilia [10]. Here we identified NBCn1 as a bona fide ciliary protein that is robustly enriched at the primary cilium in a manner dependent on distinct signals in both its N⍰and C⍰terminal cytosolic regions. Specifically, through systematic analysis of NBCn1 deletion mutants we found that amino acids 99-612 and 1118-1133 in the N- and C-terminus, respectively, must act in concert to promote NBCn1 trafficking to the cilium. In addition, we found that ciliary localization of NBCn1 is regulated by DLG1 and the retrograde IFT machinery, consistent with NBCn1 interacting physically with DLG1 [10] and the dynein 2 subunit DYNLL1 (this study). However, while DLG1 promotes trafficking of the retromer-associated protein SDCCAG3, IFT20, and polycystin-2 to primary cilia of polarized kidney epithelial cells [11], it has the opposite effect on NBCn1, which was enriched in cilia of *Dlg1*^-/-^ cells relative to WT cells. Given that NBCn1 and DLG1 are both concentrated at the basolateral membrane [7, 10, 41], which represents a way station for membrane-associated proteins destined to the primary cilium [11, 42], we propose that DLG1 may tether NBCn1 at the basolateral membrane, counteracting its endocytosis and vesicular trafficking to the cilium. In *Dlg1*^-/-^ cells, NBCn1 is released from the basolateral membrane through the loss of this anchoring, thereby permitting vesicular transport of NBCn1 to the primary cilium. This pathway may involve components of the retromer and WASH complex, which also associate with NBCn1 [7, 10] and are implicated in vesicular trafficking to the cilium [43–45].

DLG1 might also promote ciliary export of NBCn1, e.g. by facilitating its interaction with the BBSome, which functions as a membrane cargo adapter for the retrograde IFT machinery [19, 46–49]. Consistent with this model, DLG1 accumulated in BBSome-deficient mouse photoreceptor outer segments, which are modified primary cilia, supporting its link to the BBSome [50]. On the other hand, loss of IFT27, which is thought to link the BBSome with retrograde IFT trains, only caused a moderate increase in ciliary NBCn1 levels whereas inhibition of retrograde IFT with Ciliobrevin D led to robust and significant ciliary accumulation of NBCn1. These observations suggest that NBCn1 relies on retrograde IFT for ciliary export and may be associated with retrograde IFT trains independently of the BBSome. Strikingly, AlphaFold3 structural modeling and immunoprecipitation analysis identified an N⍰terminal binding motif in NBCn1 that interacts directly with the dynein light chain DYNLL1, a key component of the dynein 2 complex [27, 51–53], suggesting that ciliary export of NBCn1 is mediated by direct interaction with DYNLL1. Notably, the predicted DYNNL1 binding site in NBCn1 (residues 44–54) does not overlap with the regions required for ciliary targeting (residues 99-612 and 1118-1133) supporting the idea that DYNLL1 facilitates export of NBCn1 from cilia rather than its targeting to the organelle, consistent with our Ciliobrevin D data.

Our AlphaFold3 structural modeling and immunoprecipitation analysis also identified an interaction of residues 44–54 of NBCn1 with the endomembrane trafficking regulator VPS45. Interestingly, VPS45 has been shown to regulate ciliary homeostasis of another key ciliary ion transport protein, polycystin-2, in *C. elegans* sensory neurons by promoting its endocytic retrieval from the periciliary membrane compartment [28]. These observations support that shared mechanisms are involved in regulating ciliary homeostasis of polycystin-2 and NBCn1, and that VPS45 could function downstream of DYNLL1 and retrograde IFT to mediate endocytic retrieval of NBCn1 from the periciliary membrane compartment. Further experiments are required to test this idea.

### NBCn1 activity is dispensable for ciliogenesis but essential for ciliary length control and maintenance

Genetic ablation of *Slc4a7* in IMCD3 cells impaired bicarbonate⍰dependent pH□ recovery confirming loss of transporter function. These cells also exhibited substantially shorter cilia compared to controls, and pharmacological inhibition of NBCn1 with S0859 phenocopied this defect when applied during cilium assembly, but not when added after cilia were fully assembled, indicating a role for NBCn1 in cilia elongation. Furthermore, neither genetic ablation of *Slc4a7* nor S0859 treatment measurably altered the frequency of ciliation, supporting that NBCn1 is not required for initiation of ciliogenesis. Moreover, NBCn1 loss did not change the kinetics of serum⍰induced deciliation, as ciliation frequency and disassembly rates during serum re⍰addition in *Slc4a7^-/-^* cells were indistinguishable from those in WT. These data argue that NBCn1 is not a core initiator of ciliogenesis or ciliary resorption; rather, its activity sculpts ciliary architecture once a cilium exists. Given that ciliary length and shape are exquisitely sensitive to ionic composition and pH [31, 32], we propose that NBCn1 tunes a local bicarbonate and/or pH microenvironment that supports axonemal and membrane processes required for ciliary elongation, without directly governing ciliogenesis onset or resorption. In particular, since depletion of DYNLL1 was shown to cause reduced ciliary length in different murine cell types [54], we speculate that NBCn1 may affect ciliary length through modulation of DYNLL1 localization or activity, although this remains to be clarified. In this regard it is interesting to note that DYNLL1 dimerization has been found to be pH sensitive, by virtue of a key histidine residue in the dimer interface [55, 56].

### NBCn1 regulates Shh signaling by controlling ciliary SUFU localization

A key discovery in this study is that NBCn1 is a modulator of the Shh pathway. Whereas SMO agonists elicited the expected strong *Gli1* transcription in WT cells, this was nearly ablated in NBCn1⍰null cells. Direct stimulation with Shh ligand similarly produced attenuated *Gli1* expression in NBCn1⍰null cells compared to controls, confirming that this phenotype extends to ligand⍰driven Shh pathway activation. Mechanistically, NBCn1 deficiency did not affect ciliary dynamics of GFP⍰SMO, indicating that the earliest step of the Shh pathway is intact, but led to SUFU enrichment throughout the primary cilium in the basal state. Ordinarily, SUFU accumulates, along with GLI transcription factors, at the distal tip of the cilium in response to Shh pathway activation [37, 57, 58]. Here, we propose that NBCn1 deficiency causes SUFU ciliary accumulation at basal level, which in turn leads to dysregulation of the pathway following Shh pathway activation. While it remains to be determined how NBCn1 interacts with SUFU to control its ciliary localization, one possibility is that it acts via DYNLL1 to facilitate IFT-mediated export of SUFU from cilia during the basal state. Another possibility is that NBCn1 regulates SUFU through TMEM216, which we found interacted with NBCn1 in immunoprecipitation assays. Indeed, TMEM216, a ciliary transition zone protein [59, 60], has been reported to actively modulate Shh pathway activation via its interaction with SUFU [40].

### A working model linking ciliary NBCn1 trafficking and Shh signaling

Integrating our observations, we propose the following model. (i) NBCn1 is trafficked in vesicles from the basolateral membrane to the ciliary compartment, and this requires distinct ciliary targeting motifs in the N-terminal (residues 99-612) and C-terminal (residues 1118-1133) cytosolic regions of NBCn1 (Fig. 4A, B), which are similar but not identical to those required for its basolateral localization [10]. (ii) Following entrance to the ciliary compartment, NBCn1 is exported from the organelle by retrograde IFT, facilitated by direct interaction with DYNLL1 that binds residues 44-54 of NBCn1 (Fig. 4A, B). (iii) After exiting the cilium, NBCn1 is handed over from DYNLL1 to VPS45, which binds to the same region in NBCn1 and facilitates its endocytic retrieval from the periciliary membrane compartment. Within the cilium, NBCn1 establishes a local bicarbonate⍰rich, pH⍰buffered microdomain that supports axonemal extension, e.g. by modulating DYNLL1 or TMEM216 localization or activity. This is also required for regulating ciliary SUFU localization, ensuring proper Shh pathway activity in response to agonist- or ligand stimulation that culminates in downstream target gene (e.g. *Gli1*) expression (Fig. 4B). This model is consistent with our findings that NBCn1 loss shortens cilia and dampens Shh signaling yet leaves ciliation frequency and deciliation kinetics largely unaffected.

**Figure 4:**
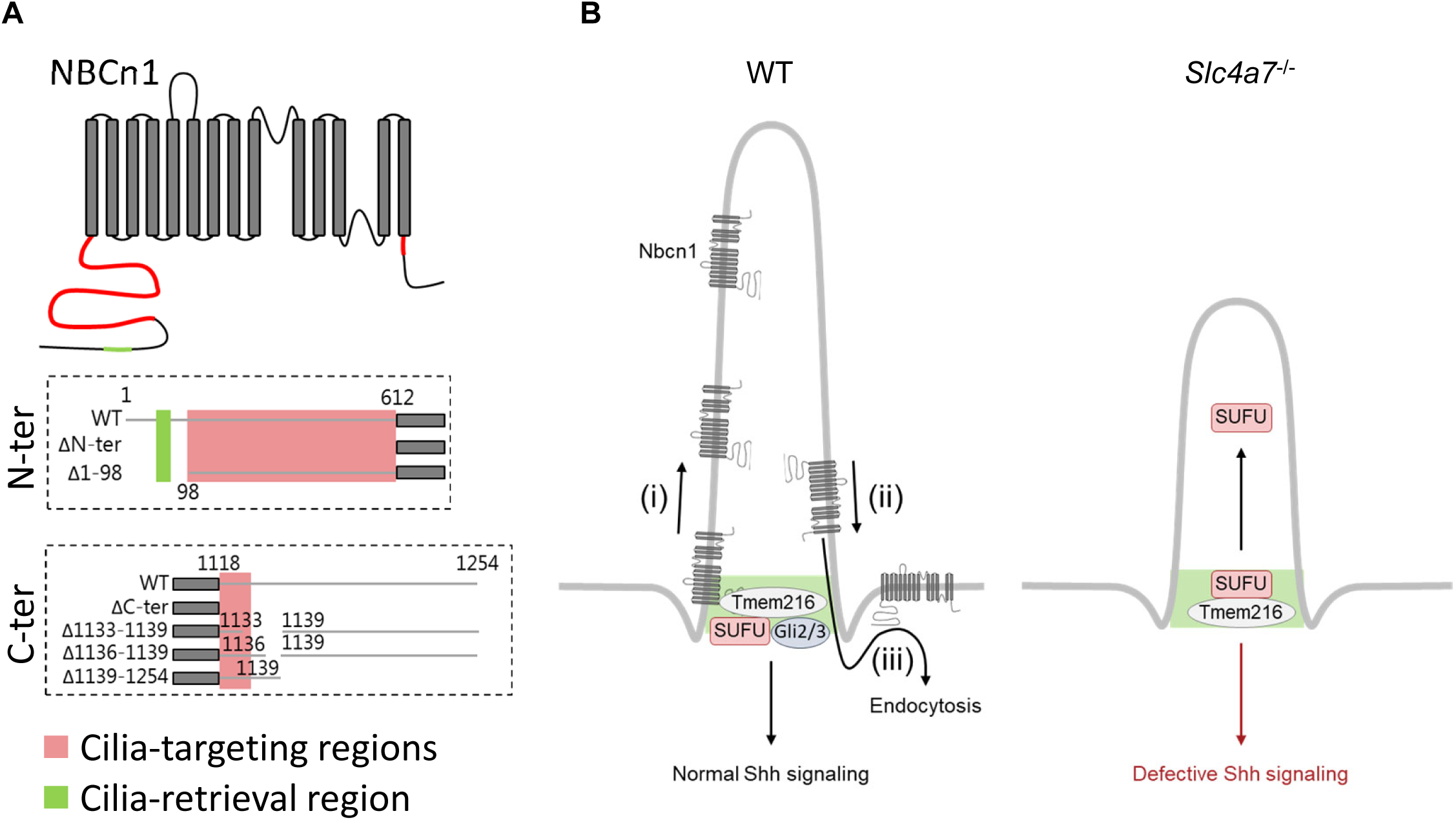
Concluding model of NBCn1 ciliary targeting and function in Hedgehog signaling. **A.** Schematic representation of NBCn1 highlighting the N- and C-terminal domains required for ciliary targeting and retrieval. **B.** Schematic representation showing the proposed mechanism by which NBCn1 is targeted and retrieved from the primary cilium: (i) NBCn1 is trafficked in vesicles from the basolateral membrane to the ciliary compartment, and this requires distinct ciliary targeting motifs in the N-C-terminal cytosolic regions (shown in A); (ii) NBCn1 is exported from the cilium by retrograde IFT, facilitated by direct interaction with DYNLL1; (iii) after exiting the cilium, NBCn1 is handed over to VPS45, which facilitates its endocytic retrieval from the periciliary membrane compartment. Furthermore, NBCn1 interferes with the interaction between DYNLL1 or TMEM216 and SUFU, thereby affecting the ciliary targeting of SUFU in *Slc4a7*^-/-^ cells, leading to defective Shh signaling.

## Conclusion

In sum, NBCn1 possesses distinct domains within its N⍰and C⍰terminal cytosolic regions that target it to the primary cilium. DLG1 and the retrograde IFT machinery negatively regulate ciliary NBCn1 levels, likely by counteracting its trafficking to (DLG1) or mediating its export from (retrograde IFT) the organelle. The latter process involves direct binding of NBCn1 to DYNLL1, which hands over NBCn1 to VPS45 following export from the cilium. Finally, NBCn1 activity is required for proper ciliary elongation and Shh signaling, owing to its interactions with and possible regulation of DYNLL1 and TMEM216 ciliary localization or function. This in turn leads to proper regulation of ciliary accumulation of SUFU in response to Shh agonist or ligand stimulation.

Our pH_i_ measurements confirm a deficit in bicarbonate⍰dependent pHi recovery in *Slc4a7* mutants at the whole⍰cell level, but future studies using cilia-targeted pH reporters are needed to resolve ciliary versus cytosolic pH microdomains and to test whether local pH balance is the operative signal by which NBCn1 controls ciliary length and SUFU localization. Given the known key roles of NBCn1 in major human disorders such as hypertension and cancer [4–6] and that dysregulated ciliogenesis and impaired Shh signaling underlie a wide spectrum of human disorders, from congenital malformations and ciliopathies to cancers driven by aberrant GLI activation, understanding how NBCn1 governs ciliary pH homeostasis may help identify new mechanistic links between ion transport dysfunction and disease.

## Materials and Methods

All reagents and solutions are listed in Table 1 and Table 2.

**Table 1.**
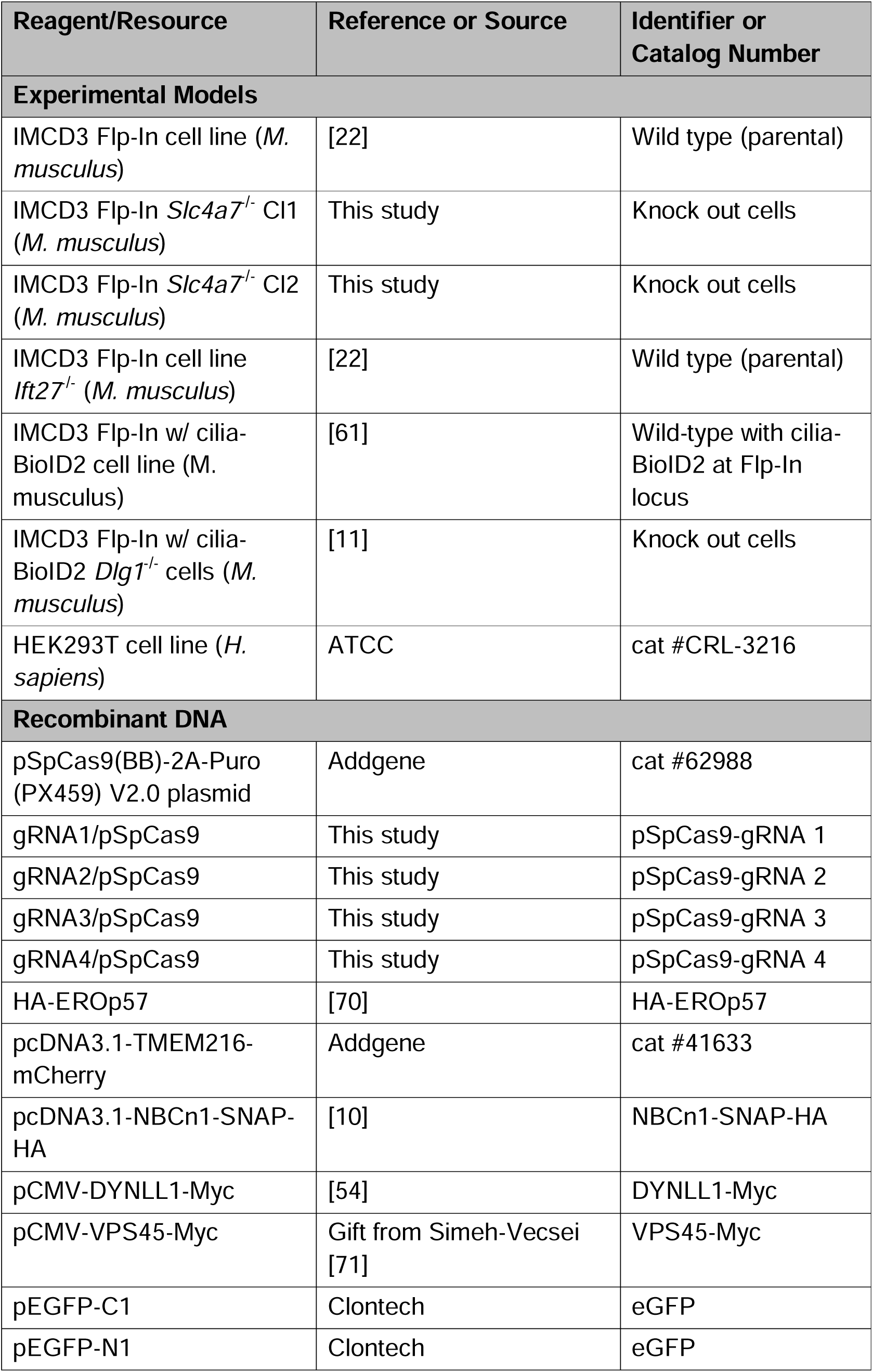
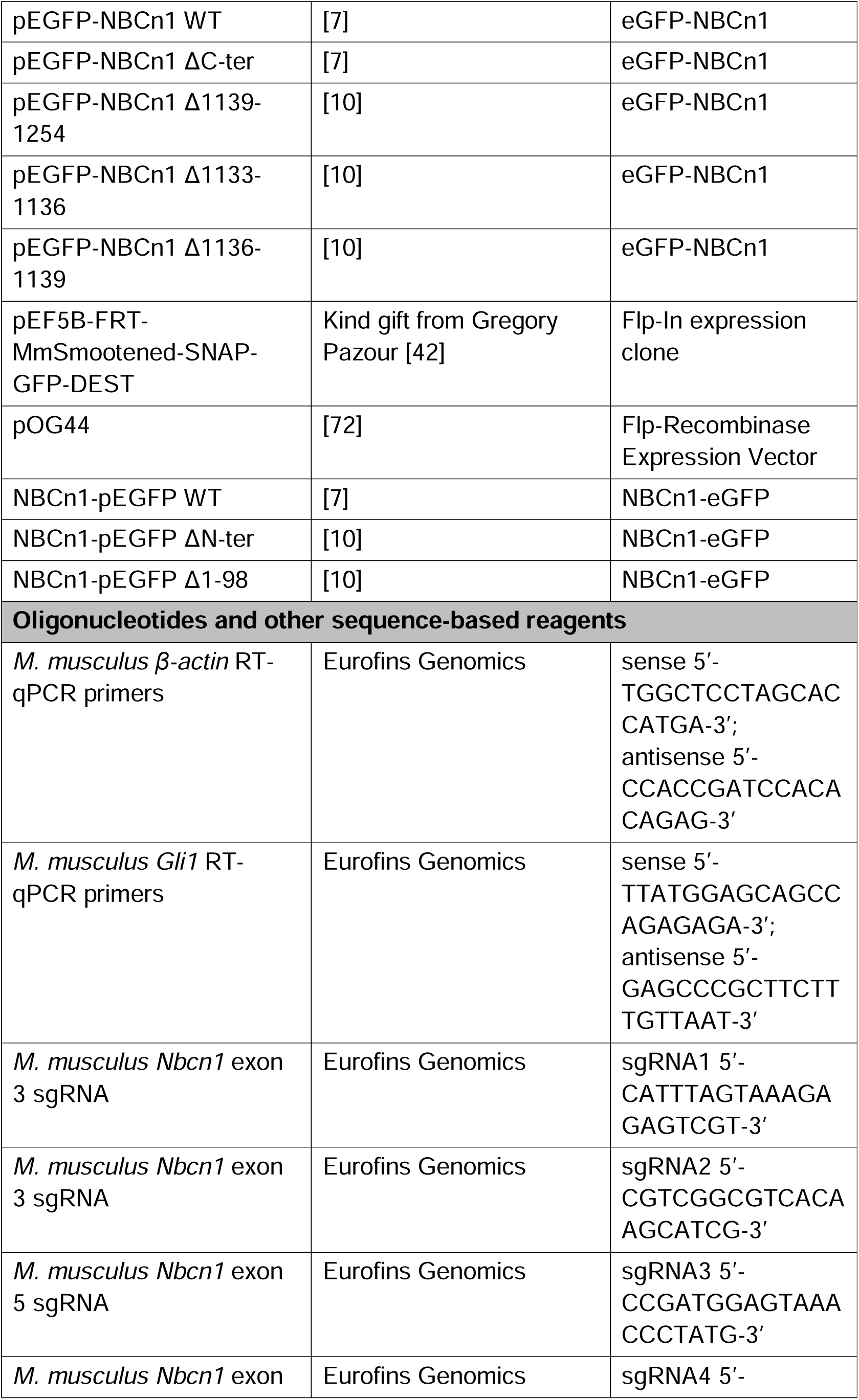
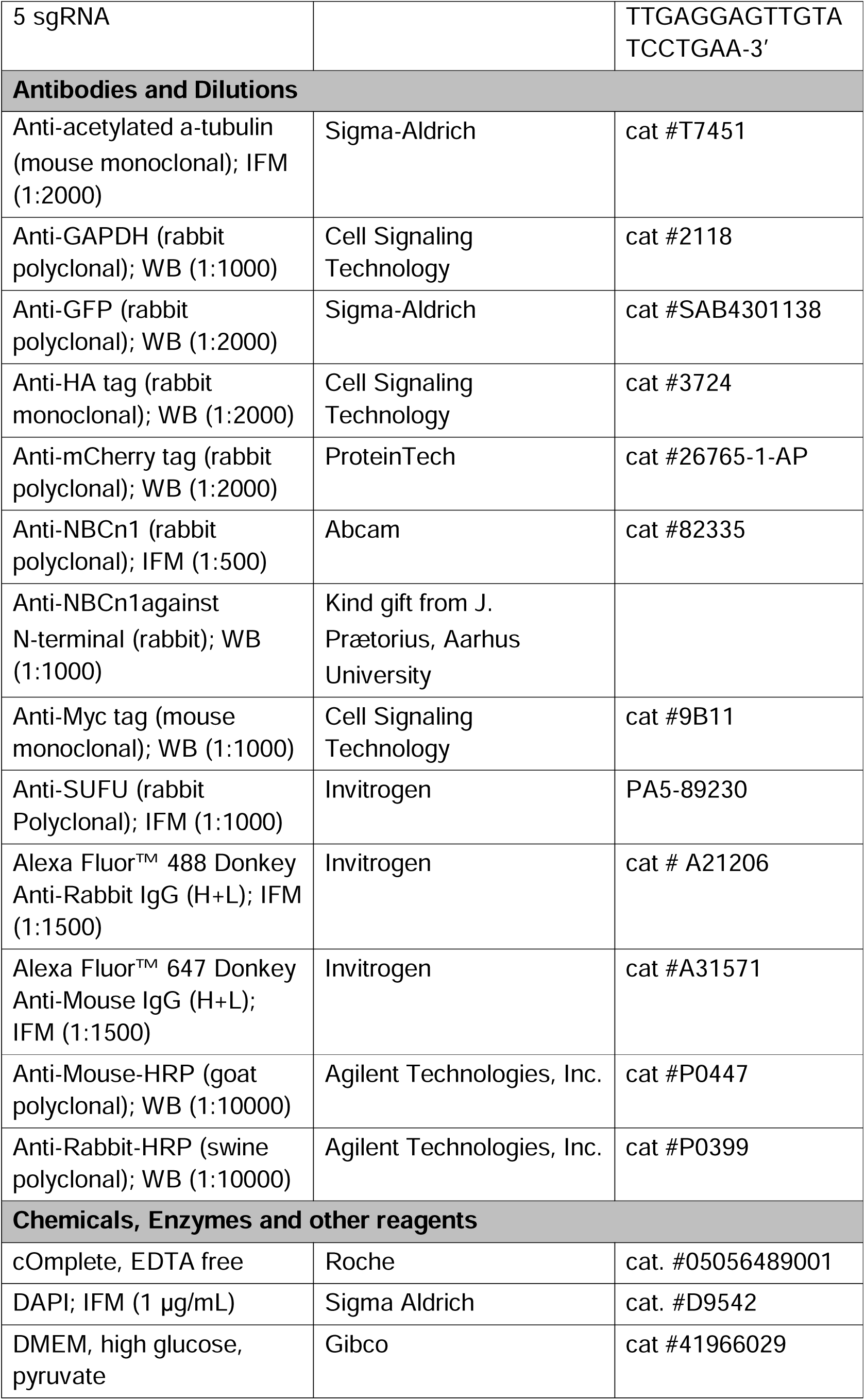
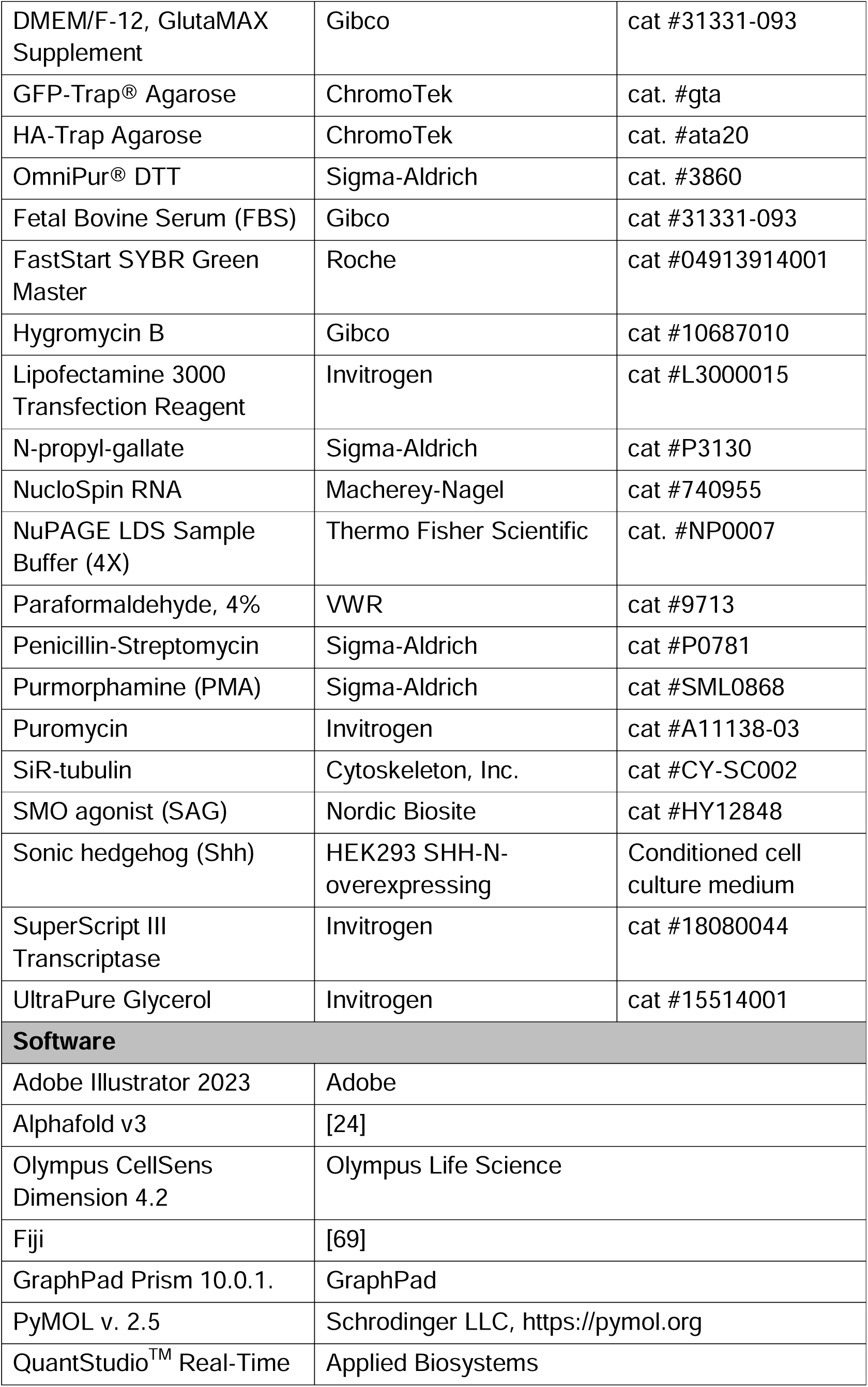
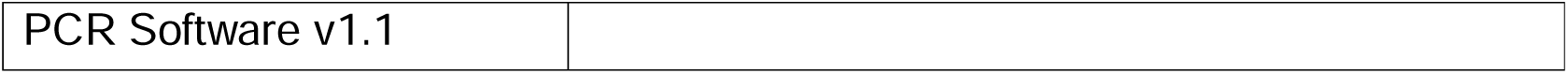

**Table 2.**
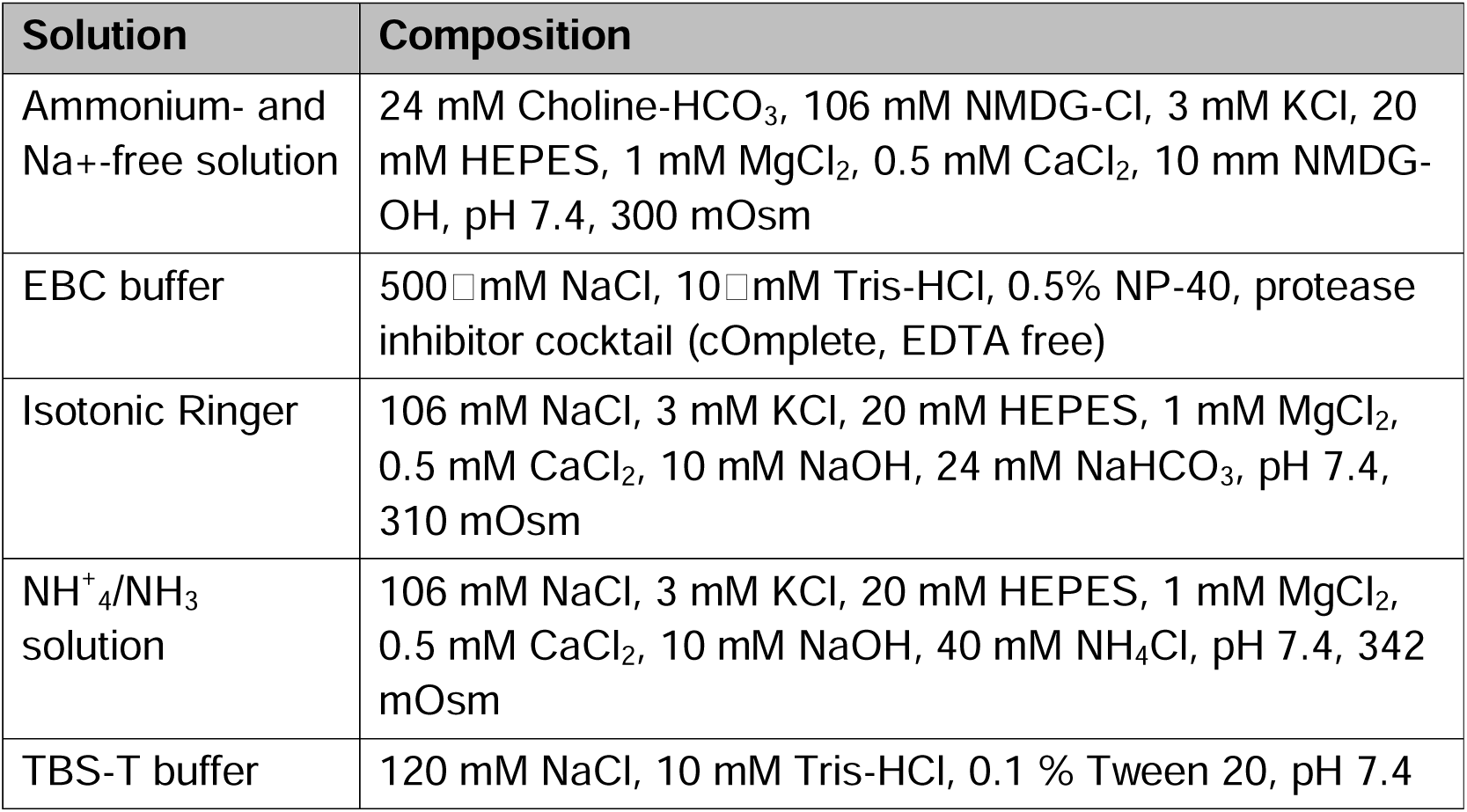

### Cell lines and reagents

An overview of reagents and cell lines described in this work is shown in Table 1. WT and *Ift27*^-/-^ IMCD3 Flp-In cell lines were a kind gift from Dr. Maxence Nachury from the University of California, San Francisco (UCSF), USA [22]. WT IMCD3 were from [61] and *Dlg1*^-/-^ IMCD3 cells were previously produced in the lab [11]. Human embryonic kidney (HEK) 293 cells were from ATCC.

### Cell culture

WT, *Ift27*^-/-^, *Dlg1*^-/-^, and *Slc4a7*^-/-^ IMCD3 Flp-In cells were cultured in DMEM/F-12, supplemented with GlutaMAX, 10% fetal bovine serum (FBS) and 1% Penicillin–Streptomycin (P/S). HEK293 cells were cultured in high-glucose DMEM supplemented with 10% FBS and 1% P/S. All cell lines were grown in a 95% humidified incubator at 37□°C with 5% CO_2_ and were routinely tested for *Mycoplasma* contamination by standard approaches. To induce ciliogenesis, IMCD3 cells were grown in absence of FBS for 24□h. Inhibition of dynein 1 and 2 or NBCn1 was followed by 24 h in this medium with 10 µM Ciliobrevin D or 20 µM S0859. To activate Shh signaling, the initial 24 h in FBS-free medium was followed by 24 h incubation with 2 µM PMA or 250 nM SAG or Shh [62] in FBS-free medium.

### Generation of NBCn1 knockout kidney epithelial cell lines

To knock out *Slc4a7* in IMCD3 cells, we employed CRISPR/Cas9 technology and used four sgRNA sequences (Table 1) from the Wellcome Sanger Institute Genome Editing tool [63]. The sgRNA spacers were cloned into the pSpCas9(BB)-2A-Puro (PX459) V2.0 plasmid as in [64]. Following phosphorylation and annealing of the two complementary sgRNA oligos, they were ligated into the BbsI-digested backbone. Selected clones were sequenced to verify the spacer sequence. WT IMCD3 cells were transfected with the four pooled Cas9-gRNA plasmids (using Lipofectamine 3000. A day after transfection, cells were treated with 2□µg/mL puromycin for five days and tested for NBCn1 depletion by western blot analysis. Selected clones were validated by western blot analysis, Sanger sequencing, and pH_i_ recovery analysis.

### Generation of transgenic cell lines

WT and *Slc4a7*^-/-^ IMCD3 Flp-In cells were stably transfected to express SMO using the Thermo Fisher Scientific Flp-In™ technology. Briefly, cells were transfected with equal amounts of plasmid DNA encoding pEF5B-FRT-MmSmootened-SNAP-GFP-DEST [42] and the pOG44 Flp-In recombinase plasmid DNA by lipofection using Lipofectamine 2000, followed by clonal selection with 400 ug/ml Hygromycin B.

### Measurements of pH_i_

WT and *Slc4a7*^-/-^ IMCD3 cells, grown in quintuplicate in 48 well plates, were loaded with the ratiometric pH indicator BCECF by incubation with 1.6 μM BCECF-AM in FBS-free medium for 30 min, wash and 30 min rest. Cells were switched to isotonic HCO_3_^-^ containing Ringer (Table 2) and plates were placed at 37°C/5% CO_2_ in a Varioskan™ LUX Multimode Microplate Reader (Thermo Scientific). pH_i_ was determined by ratiometric analysis of emission at 530 nm after excitation at 490 and 440 nm excitation. After determining base-line pH_i_, cells were alkalinized with an NH_4_^+^/NH_3_ solution (Table 2), which was replaced by Na^+^-free solution (Table 2) to induce rapid acidification, and finally with isotonic Ringer to monitor Na^+^-dependent pH_i_ recovery. The difference between the initial rate of pH_i_ recovery in WT and *Slc4a7*^-/-^ IMCD3 cells reflects the loss of NBCn1 activity in the KO cells.

### Alpha Fold modeling of protein complexes

AlphaFold3 [24] was used to model pairwise interactions between full-length mouse NBCn1 (UniProt Q8BTY2, 1034 aa) and candidate interaction partners from the human ciliary proteome [25, 65]. Predictions were performed using AlphaFold3 v3.0 with default parameters, generating five independent structural samples per protein pair using different random seeds. For each prediction, the interface predicted template modeling score (iPTM) was used as the primary metric of interaction confidence, where iPTM > 0.6 indicates a plausible interaction. The predicted aligned error (PAE) was used to assess local confidence in the relative positioning of residues between chains, and the predicted local distance difference test (pLDDT) was used to evaluate per-residue model confidence. Interface residues were identified using a combined spatial filter (Cbeta-Cbeta distance < 8 Angstrom) and PAE threshold (PAE < 10 Angstrom). Structure visualization and figure preparation were performed using PyMOL (Schrodinger LLC). Structures are colored by pLDDT confidence (blue: very high, pLDDT > 90; cyan: confident, 70 < pLDDT < 90; yellow: low, 50 < pLDDT < 70; orange: very low, pLDDT < 50).

### Immunoprecipitation, SDS-PAGE and western blot analysis

Transient transfection of AlphaFold3 candidates and immunoprecipitation (IP) analysis in HEK293 cells were performed following the protocol described by [66]. Proteins were harvested on ice cold with EBC buffer. For GFP and HA IP experiments, protein extracts were incubated 2□h with either 15□μL HA-Trap Agarose or GFP-Trap Agarose beads. Samples from IP analysis were prepared for SDS-PAGE analysis by addition of NuPAGE LDS Sample Buffer and 50 mM DTT. Input and immunoprecipitated fractions were analyzed by SDS-PAGE and western blotting as outlined in [66], using the antibodies and dilutions listed in Table 1. SDS-PAGE and western blotting of IMCD3 cell lysates were performed similarly.

### Immunofluorescence microscopy

IMCD3 cells were grown on coverslips to 60-70% confluency, followed by 24 h culture in absence of FBS (unless stated otherwise). Cells were fixed in 4% paraformaldehyde for 15 min, permeabilized in 1% bovine serum albumin (BSA) and 0.2 % Triton X-100 in phosphate-buffered saline (PBS) for 10 min, and blocked in 5% BSA in PBS for 1 h, all at room temperature. Cells were incubated with primary antibodies in 1% BSA in PBS overnight at 4°C, washed 3 times in TBS-T, and incubated 1 h in secondary antibodies in 1% BSA in PBS, for 1 h at room temperature. Nuclei were stained with DAPI, and coverslips were finally mounted on a slide with 2% n-propyl gallate diluted in UltraPure Glycerol and sealed. Slides were imaged using an Olympus inverted microscope (IX83) equipped with a Yokogawa spinning disc unit and a Teledyne Prime 95B back-illuminated sCMOS camera (01- PRIME-95B-R-M-16-C), using the 100× oil objective. The 405 nm, 488 nm and 640 nm laser lines were used to image DAPI, Alexa Fluor™ 488 and Alexa Fluor™ 647 antibodies, respectively.

### Time-lapse live cell imaging

WT and mutant IMCD3 cells were seeded in 35 mm glass-bottom dishes, serum-starved for 24 h, and transfected for 6 h with 2 μg pEGFP-NBCn1 or NBCn1-pEGFP (WT or mutants) using Lipofectamine-3000. To visualize primary cilia, culture medium was supplemented with 1 μM SiR-tubulin for at least 45 min prior to imaging. Dishes were mounted in the environmentally controlled chamber (37°C/5% CO_2_) of an Olympus inverted microscope (IX83) equipped with a Yokogawa spinning disc unit and a Teledyne Prime 95B back-illuminated sCMOS camera (01-PRIME-95B-R-M-16-C), and cells were imaged every 3 s using the 100× oil objective. The 488 nm and 640 nm laser lines were used to image GFP-NBCn1 and SiR-Tubulin-Cy5, respectively.

### Quantitative RT-PCR

Total RNA was isolated using NucleoSpin RNA and concentration and purity were determined using a Biodrop Duo spectrophotometer. 1 μg RNA was reverse transcribed using SuperScript III Transcriptase to generate cDNA, which was then amplified in triplicates using a SYBR Green and primer mastermix in a QuantStudio 7 Flex Real-Time PCR System with the following steps: 95°C for 10 min; 40 cycles of (95°C for 30 s, 57°C for 1 min, 72°C for 30 s); then 95°C for 1 min. Primers were designed using Primer3Plus (https://www.primer3plus.com), synthesized by Eurofins Genomics and diluted in nuclease-free H_2_O. mRNA levels were determined using the Pfaffl method [67], with β-actin as housekeeping gene.

### Quantitative and statistical analysis of IFM and western blot data

IFM data were processed and analyzed using methods previously described [11, 68]. Ciliary length and frequency, and relative mean fluorescence intensity (MFI) measurements of relevant antibody-labeled antigens within the cilia in WT and mutant cells. For MFI analysis, the background-corrected MFI was normalized to relevant control cells. For quantitative western blot analysis, the average pixel intensity of the bands was measured using Fiji software [69] and normalized against loading control. All quantitative data are presented as mean□±□standard error of the mean (SEM). Statistical analysis was performed using GraphPad Prism 10, with the specific test indicated in the figure legends. Significance levels are indicated in the figure legends. Unless otherwise noted, all experiments were performed in at least three independent biological replicates.

## Acknowledgements

Supported by grants from the Novo Nordisk Foundation (NNF22OC0080406) and the Carlsberg Foundation (CF22-0670). We thank Maxence Nachury for *Ift27* mutant cells.

## Figure legends

<u>Figure S1</u>: **A.** Confocal images of WT and *Dlg1*^-/-^ cells expressing GFP-tagged NBCn1 full-length/wild-type (WT). SiR-tubulin (magenta) stains primary cilia. Insets show magnified views of the cilium (“shifted” panels). Scale bar, 10 µm. **B.** IFM analysis of ciliated IMCD3 cell lines showing comparative ciliary GFP-NBCn1 staining in WT and *Dlg1*^-/-^ cells. Each red point represents one biological replicate; grey violin plots represent the technical replicates; bars indicate mean ± SEM (N= 3, WT n= 124, *Dlg1*^-/-^ n= 104) **C.** Representative confocal images of WT and *Ift7*^-/-^ cells before and after treatment with 10 μM Ciliobrevin D stained for IFT88 (green), acetylated α-tubulin (magenta) to visualize primary cilia for nuclei. Scale bar, 10 µm. **D.** IFM analysis of ciliated IMCD3 cell lines showing comparative ciliary IFT88 staining in WT and *Ift27*^-/-^ cells before and after treatment with 10 μM Ciliobrevin D. Each red point represents one biological replicate; grey violin plots represent the technical replicates; bars indicate mean ± SEM (N= 3, WT DMSO n= 134, *Ift27*^-/-^DMSO n= 99, WT Ciliobrevin D n= 104, *Ift27*^-/-^ Ciliobrevin D n= 111); statistical significance was determined by two-way ANOVA with multiple comparison testing. **E.** Predicted aligned error (PAE) matrix for the NBCn1–DYNLL1 complex (iPTM = 0.64). The interface comprises 11 contacts with PAE < 6 Å and 47 contacts with PAE < 10 Å. Dark green indicates high confidence in relative residue positioning. **F.** Predicted aligned error (PAE) matrix for the NBCn1–VPS45 complex (iPTM = 0.61). The interface comprises 7 contacts with PAE < 6 Å and 25 contacts with PAE < 10 Å.

<u>Figure S2</u>: **A, B.** Quantification of cilia frequency in WT and *Slc4a7*^-/-^ clones at 24 h (A) and 48 h (B) post FBS starvation. Each black/red point represents one biological replicate; Grey violin plots represent the technical replicates; bars indicate mean ± SEM (N= 3; 24 h WT n= 18, 24 h *Slc4a7*^-/-^ Cl1 n= 14, 24 h *Slc4a7*^-/-^ Cl2 n= 13, 48 h WT n= 13, 48 h *Slc4a7*^-/-^ Cl1 n= 13, 48 h *Slc4a7*^-/-^ Cl2 n= 13); statistical significance was determined by one-way ANOVA with multiple comparison testing. **C.** Quantification of cilia frequency after S0859 treatment. Cilia frequency was measured following S0859 treatment before (24h) or after (24 h + 24 h) ciliogenesis. Each red point represents one biological replicate; grey violin plots represent the technical replicates; bars indicate mean ± SEM (N= 3; 24 h DMSO n= 12, 24 h S0859 n= 11, 24 h + 24 h DMSO n= 12, 24 h + 24 h S0859 n= 11); statistical significance was determined by two-way ANOVA with multiple comparison testing.

<u>Figure S3</u>: **A.** Experimental timeline for analysis of FBS-induced deciliation. **B.** Representative confocal images of WT and *Slc4a7*^-/-^ cells (Cl1, Cl2) stained for acetylated α-tubulin (magenta) to visualize primary cilia and DAPI (blue) for nuclei. Scale bar, 10 µm. **C.** Quantification of cilia frequency in WT and *Slc4a7*^-/-^ clones at 0, 1, 2, 3, 4, 5 and 6 h post FBS stimulation, as well as 24 h starvation post FBS stimulation. Each black/red point represents one biological replicate; bars indicate mean ± SEM (N= 3); statistical significance was determined by one-way ANOVA with multiple comparison testing. **D.** Quantification of cilia disassembly rate in WT and *Slc4a7*^-/-^ clones after FBS stimulation. Each black/red point represents one biological replicate; bars indicate mean ± SEM (N= 3); statistical significance was determined by one-way ANOVA with multiple comparison testing.

<u>Figure S4</u>: **A.** RT-qPCR of *Gli1* expression in WT and *Slc4a7*^-/-^ cells following stimulation with Shh. Each black/red point represents one biological replicate; bars indicate mean ± SEM (N= 3); statistical significance was determined by two-way ANOVA with multiple comparison testing. **B.** Confocal images of cells expressing GFP-SMO. Scale bar, 10 µm. **C**-**E.** IFM analysis of ciliated IMCD3 cell lines showing comparative SMO ciliary localization upon PMA and SAG treatment in WT (C) and *Slc4a7*^-/-^ (D, E) cells. Each red point represents one biological replicate; grey violin plots represent the technical replicates; bars indicate mean ± SEM (N= 3; DMSO WT n= 58, DMSO *Slc4a7*^-/-^ Cl1 n= 83, DMSO *Slc4a7*^-/-^ Cl2 n= 72, PMA WT n= 58, PMA *Slc4a7*^-/-^ Cl1 n= 83, PMA *Slc4a7*^-/-^ Cl2 n= 73, SAG WT n= 72, SAG *Slc4a7*^-/-^ Cl1 n= 58, SAG *Slc4a7*^-/-^ Cl2 n= 88); statistical significance was determined by two-way ANOVA with multiple comparison testing.

