## Supplementary figures and images for "NBCn1 interacts with DYNLL1 and regulates ciliary length and SUFU localization to control Sonic hedgehog signaling"

### Supplemental Figures

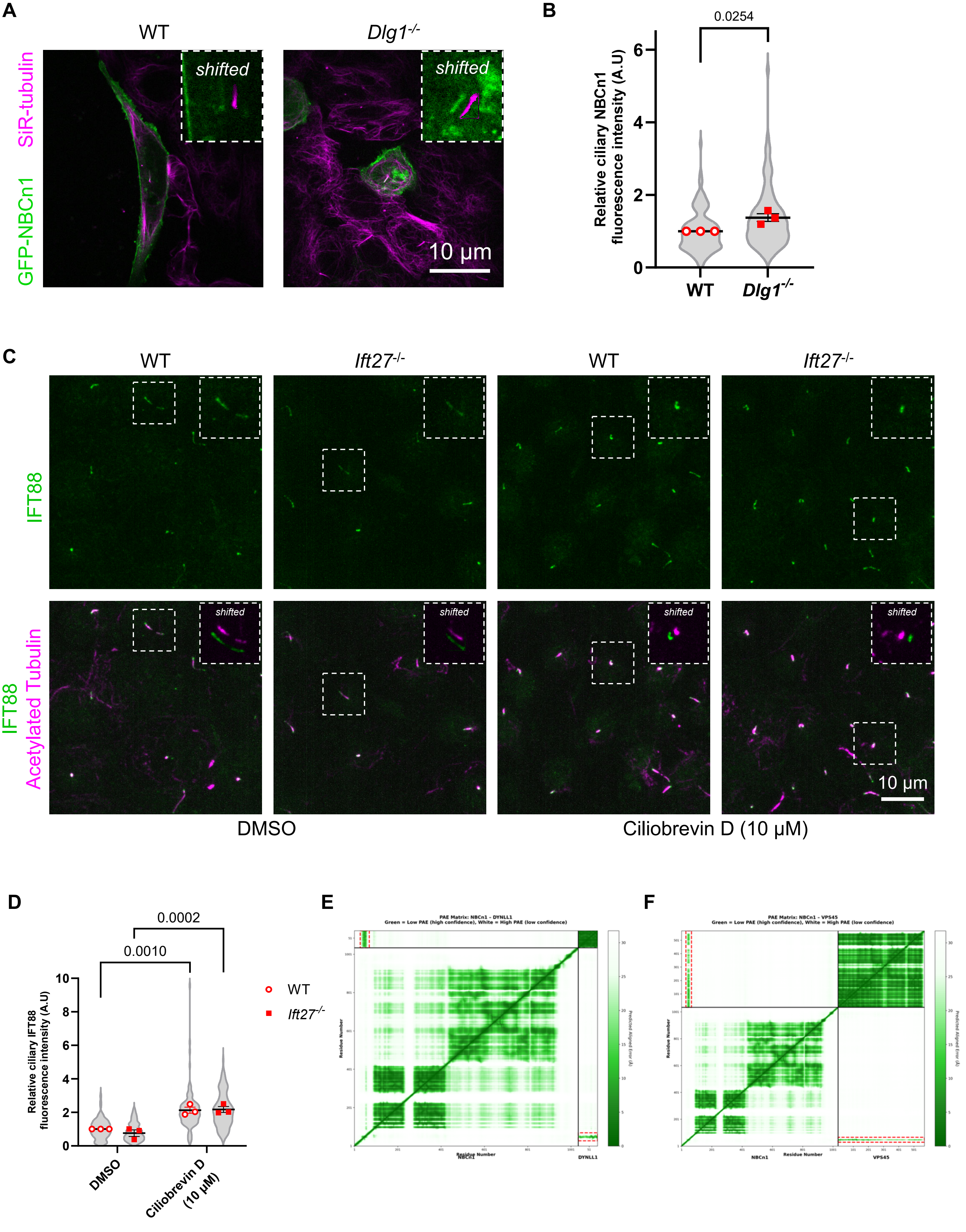

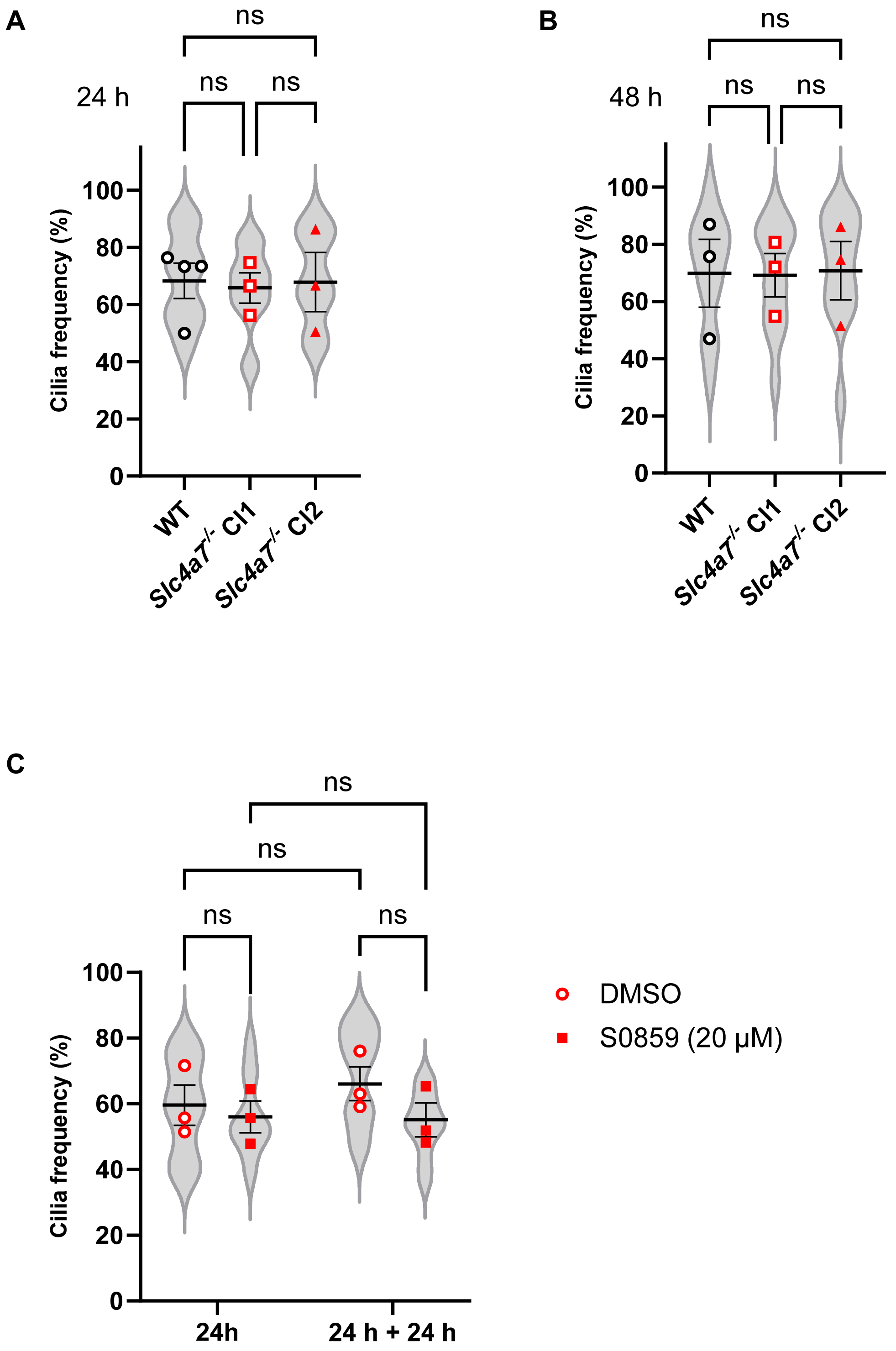

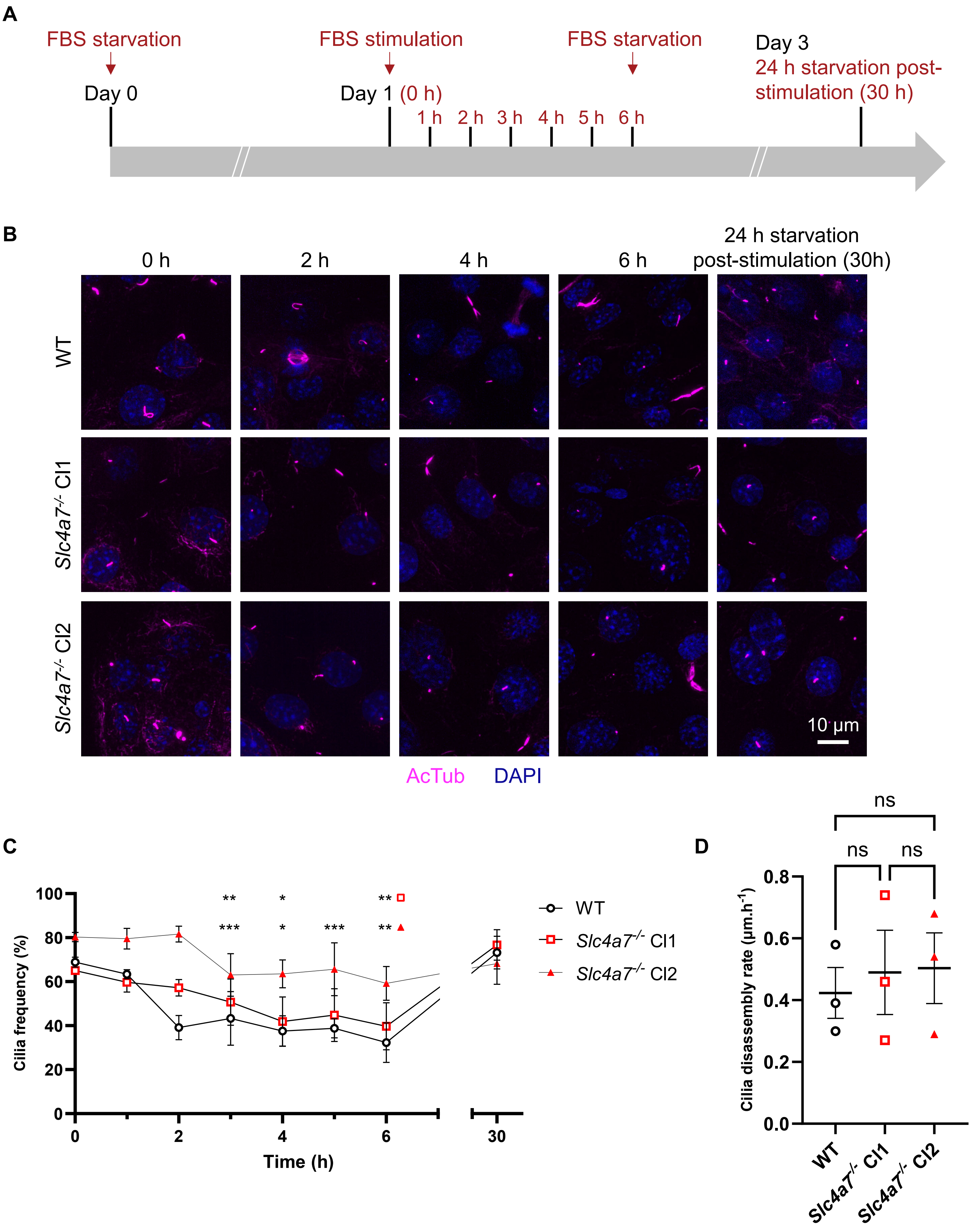

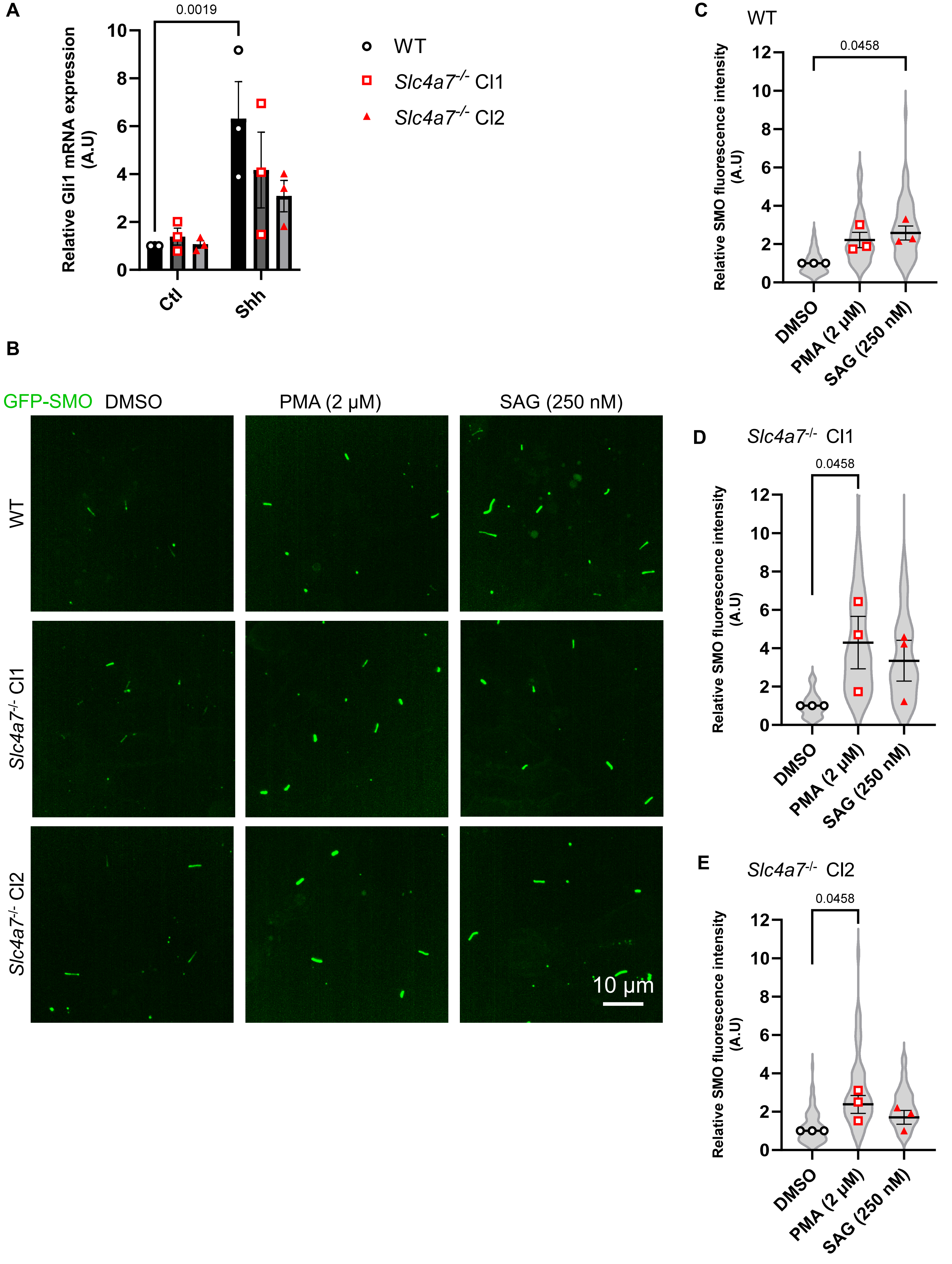
